# Linker stretching optimizes bonding entropy in sticker-spacer condensates

**DOI:** 10.64898/2026.10.09.758027

**Authors:** Sk Ashif Akram, Junlang Liu, Aniruddha Chattaraj, Sanjeev Gautam, Buddhapriya Chakrabarti, Eugene I. Shakhnovich, Jeremy D. Schmit

## Abstract

Phase separating biopolymers are often described by the sticker-spacer model, which accounts for non-attractive linkers that connect discrete attractive moieties. “Associative polymer” theories describe these systems as two coexisting fluids of stickers and spacers, respectively. In such theories the stickers and spacers are decoupled in the sense that sticker binding does not affect spacer conformations or vice versa. Informed by coarse-grained simulations, we develop a phenomenological mean-field theory to determine the microscopic coupling between the sticker and spacer degrees of freedom. Accounting for linkers connecting adjacent spacers introduces free energy contributions due to conformational entropy of the spacer and the binding combinatorics of the stickers. These contributions have opposite dependencies on spacer length because longer spacers allow a greater volume to search for binding partners but incur a greater entropy cost upon bond formation. The length dependent terms from these contributions cancel leaving a length-independent elastic term arising from the fact that a small amount of spacer stretching lowers the free energy by increasing the number of possible binding partners. The cancellation of spacer length from the theory justifies the decoupling approximation and supports mean-field treatments of heterogeneous spacers.

## Introduction

Polymer phase separation drives the formation of numerous membraneless organelles within biological cells.^1–3^ In contrast to the synthetic homopolymers that motivated Flory-Huggins solution theory and microphase separation models, ^4–6^ biomolecules are heteropolymers, implying that interactions between two molecules can have a variety of attractive and repulsive forces, depending on which portions of the molecules come into contact. This heterogeneity has motivated the “stickers and spacers” model, which describes polymers with distinct attractive “sticker” moieties that are separated by non-attractive “spacers”.^7^ Semenov and Rubinstein^8^ (hereafter referred to as “SR”) developed a comprehensive theory to describe such polymers that was later adapted to model proteins^9^ and multi-component systems.^10,11^ The SR free energy is equivalent to a description of two coexisting fluids. One fluid contains the stickers and favors higher densities to maximize bond interactions and the combinatorial entropy of binding. The second fluid contains the spacers and favors lower densities to minimize repulsive (primarily excluded volume) interactions. In SR theory the only interaction between these fluids comes from the constraint that the concentrations of the sticker and spacer fluids are related by the composition of the chains. However, other theoretical models demonstrate that constraints from the spacer chains connecting the stickers strongly influence the free energy.^12,13^

In this work we employed a combination of simulation and theory to understand how the connections between stickers affect the sticker-spacer free energy. To isolate the effect of connectivity on sticker binding, we held the total number of beads and the simulation volume fixed while varying the number and spacing of stickers along each chain. The nominal polymer volume fraction was therefore identical across these systems, ensuring that the observed changes were dominated by sticker association and connectivity. We find that the linkages between adjacent stickers introduce two free energy contributions. The first comes from the fact that longer spacers have a greater conformational entropy cost when the ends are immobilized by bond formation. The second contribution comes from the fact that longer spacers allow the adjacent sticker to search a greater volume for binding partners, which leads to a greater combinatorial entropy of binding. Interestingly, the spacer length dependence cancels between these terms leaving a length-independent term that is absorbed into the fitting parameter of theories following the SR formalism. As a result, linkage constraints do not alter the sticker-fluid framework where bond formation follows the association-dissociation equilibrium of untethered reactants. However, the competition between the spacer conformational entropy and sticker combinatorial entropy has a residual effect that stretches the segments between adjacent bonds by 4.6%.

## Model

To isolate the relationship between spacer connectivity and sticker binding, we considered systems where the number of chains, the system volume, and the occupied volume fraction are held fixed. Specifically, we used volumes small enough that the system is forced into a single dense phase, such that the mixing entropy and non-specific interaction terms are nearly unchanged upon variations in the sticker/spacer composition. This allowed us to isolate the effects of connectivity on the binding free energy. Our simulations used *N*_*A*_ = 50 chains with type “A” stickers and *N*_*B*_ = 50 chains with type “B” stickers. Each chain contained *L* = 100 total beads, of which *v*_*A*_ = *v*_*B*_ were stickers able to form heterotypic A-B bonds of affinity *ϵ*

## Results

### Theory

In the SR theory^8^ the sticker-sticker interactions contribute two terms to the free energy

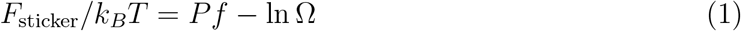

where *f* is the free energy of bond formation, expressed in units of *k*_*B*_*T, P* is the number of bonds, and Ω is the number of ways to distribute *P* bonds among the *M*_*A*_ = *N*_*A*_*v*_*A*_ type A stickers and *M*_*B*_ = *N*_*B*_*v*_*B*_ type B stickers, where *N*_*i*_ is the number of chains of type *i* and *v*_*i*_ (“valence”) is the number of stickers per chain. The combinatorial factor Ω depends on the length of spacers, which constrains the volume a particular sticker can search for a binding partner. To compute the effect of connectivity on Ω, we begin by assuming the A polymers are randomly distributed within volume *V* and count the ways to distribute the B polymers such that *P* bonds are formed. We account for connectivity by introducing a parameter, *z*, that describes the number of A stickers that are within reach of a B sticker given the constraint that the other end of the adjoining spacer is fixed in space by the previous bond (Fig. 2).^12^ With each placed bond there is an increasing probability that a potential binding site is already occupied by a bond. So, while the first sticker has *z* options to form a bond, the second sticker has *z*(*M*_*A*_ − 1)*/M*_*A*_, the third has *z*(*M*_*A*_ − 2)*/M*_*A*_, etc. This gives a total of (*z/M*_*A*_)^*P*^ *M*_*A*_!*/*(*M*_*A*_ − *P* )! options for choosing the A binding sites. This expression is equivalent to the number of conformations of a lattice polymer with *P* segments where *z* is the coordination number of the lattice (i.e., *z* = 2*d* for a *d*-dimensional square lattice). Next, we note that there are *M*_*B*_!*/*((*M*_*B*_ − *P* )!*P* !) ways to select which B sites form bonds. Thus, the number of binding possibilities is

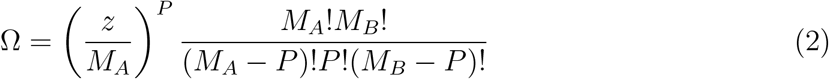

This expression is equivalent to prior work ^8,11^ except in the interpretation of the proximity term. In Eq. 2 sticker proximity is accounted for by the *z/M*_*A*_ factor, which describes the probability that a potential binding partner is within reach of the spacer. In SR theory, the equivalent factor is *v*_0_*/V* where *v*_0_ is the volume around a sticker that is within the range for bond formation. Therefore, *v*_0_*/V* is the probability that two randomly chosen stickers are close enough to form a bond.

Inserting Eq. 2 into Eq. 1 shows that the proximity term has the effect of modifying the effective sticker affinity

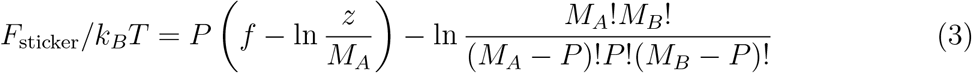

A second contribution of the spacers comes from the bond free energy *f* . Our sequential method of placing B stickers implies that each bond restricts the conformational entropy of the spacer connecting to the previous bound sticker pair. The conformational entropy can be obtained from the unconstrained end-to-end probability distribution function, which is given by

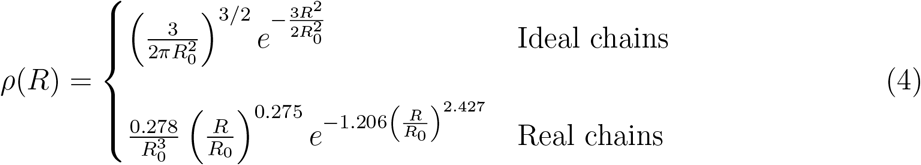

where *R* is the distance between the ends, and the distributions are normalized such that 4*π∫ ρ*(*R*)*R*^2^*dR* = 1.^14,15^ The entropic cost of constraining the end-to-end extension to a distance *R* is obtained from the logarithm of the end-to-end distribution function ^15,16^

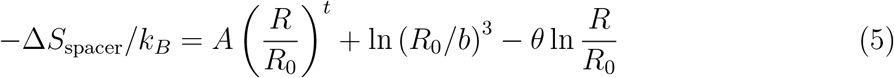

where *A* = 3*/*2, *t* = 2, and *θ* = 0 for ideal chains, and *A* = 1.206, *t* = 2.427, and *θ* = 0.275 for real chains. A microscopic length scale *b*, comparable to the Kuhn length, ^16^ has been introduced to remove dimensions from the logarithm and absorb the numerical prefactor. In Eq. 5 the first term is the elasticity, which obeys Hooke’s law in the ideal case. The remaining terms reflect the cost of choosing a specific point within the 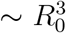 volume accessible to the polymer end, where the term proportional to *θ* corrects for the region near the origin that is inaccessible to chains with excluded volume.

The conformational entropy cost given by Eq. 5 should be included in the bond formation free energy so that *f* = *ϵ* − Δ*S*_spacer_*/k*_*B*_ where *ϵ* is the bond energy. Inserting this into the free energy we have

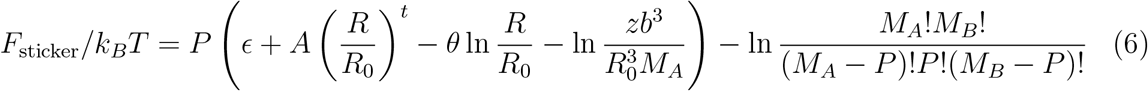

Next we use the fact that the parameter *z* describes the number of A stickers within reach of the spacer, which is the product of the sticker concentration and the volume of the accessible sphere *z* = 4*πR*^3^*M*_*A*_*/*(3*V* ) (Fig. 2). Therefore, the free energy can be written as

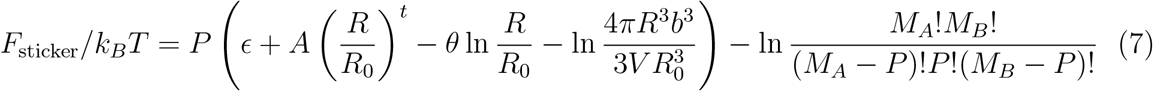

Minimizing the free energy with respect to *R* gives *R* = *R*_0_((*θ* + 3)*/At*)^1*/t*^, which evaluates to *R* = *R*_0_ for ideal chains and *R* ≃ 1.046*R*_0_ for real chains. Notably, the stretching of a bound spacer *s* = *R/R*_0_ depends only on the polymer scaling (ideal or real) and not on the spacer length. Inserting *s* into the free energy gives

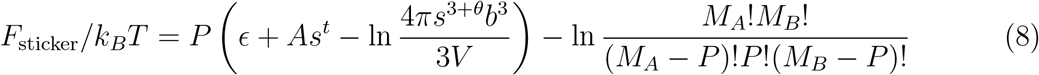

Crucially, Eq. 8 does not depend on spacer length because the length dependent quantities cancel in the *R* optimization. This cancellation is required to restore symmetry with respect to the exchange *A* ↔ *B*. The physical basis for the length cancellation is that both the spacer conformational entropy and the sticker combinatorial entropy depend on the volume accessible to a polymer that is pinned at one end. On one hand, a larger volume allows more options for the sticker at the free end to find a binding partner, which is captured by the *R*^3^ scaling of *z*. On the other hand, a larger search volume implies a greater loss of entropy when the free sticker becomes pinned to a specific binding partner, which is captured by the 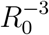 normalization constant in the polymer end-to-end distribution function. The cancellation between these length-dependent factors results in a length-independent microscopic volume 4*πs*^3^*b*^3^*/*3 divided by the system volume, which is functionally identical to the *v*_0_*/V* factor proposed by SR. This correspondence means that the entropic coupling between the sticker and spacer fluids can be absorbed by the *v*_0_ fitting parameter. Importantly, the cancellation between the search volume terms allows us to neglect variations in spacer lengths due to heterogeneities in the molecule as well as the effective doubling in the spacer length if the previous sticker not bound.

### Simulation Methods

We performed coarse-grained molecular dynamics (MD) simulations of an associative polymer system composed of two polymer types, denoted A and B, using the LAMMPS molecular dynamics package.^17^ The two chain types were chemically identical in all respects except for their labels. Each chain consisted of a total of *L* = 100 coarse-grained beads (Fig. 1). Along each chain, beads are classified as either stickers or spacers. To investigate the role of sticker density, the number of stickers per chain was varied systematically while keeping the total chain length fixed at 100 beads. Simulations were performed for uniformly distributed sticker counts of 5, 10, 20, and 25 per chain as shown in Fig. 1.

**Figure 1:**
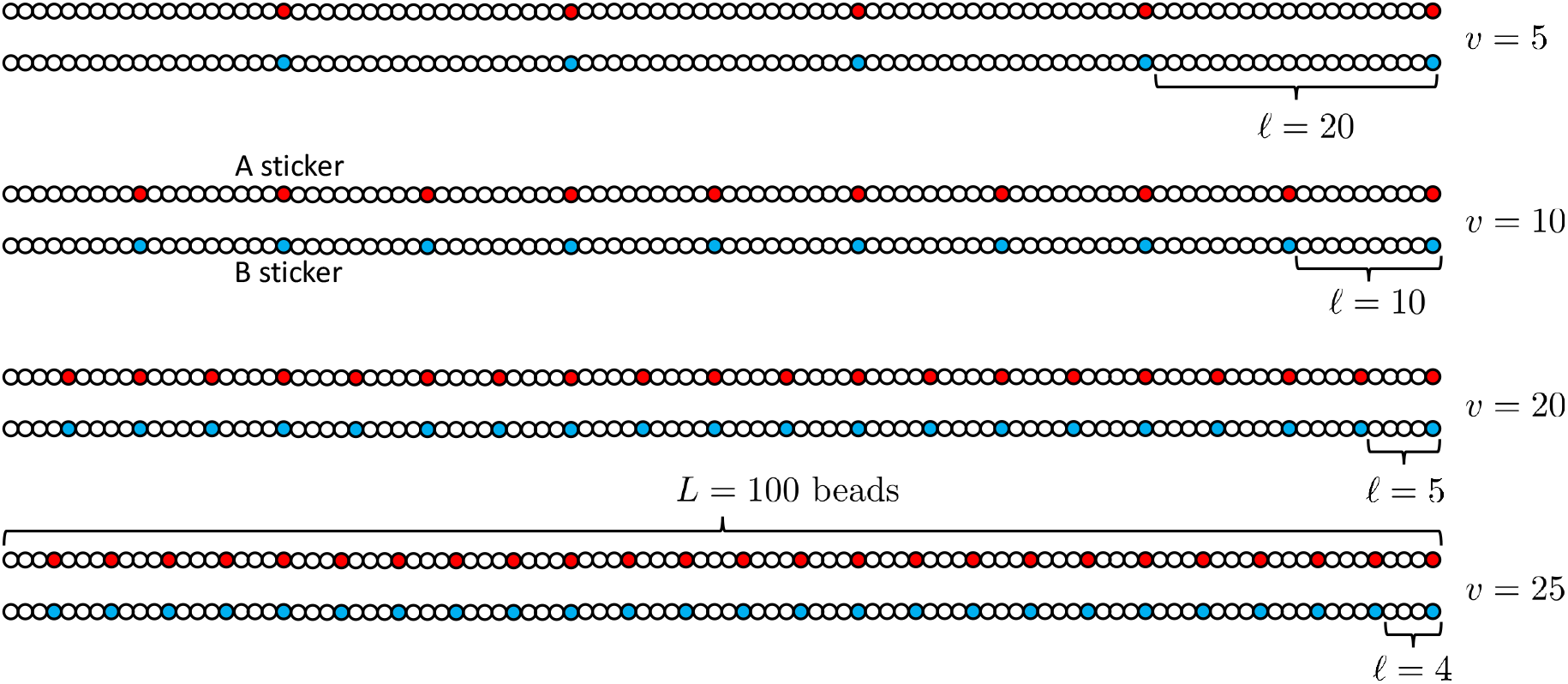
Schematic representation of chains with *L* = 100 beads, which contain *v* univalent stickers and *L*− *v* spacer beads. The spacer length *l* = *L/v* describes a contiguous stretch of spacer beads plus one adjoining sticker. Our simulations used *v* = 5, 10, 20, 25.

**Figure 2:**
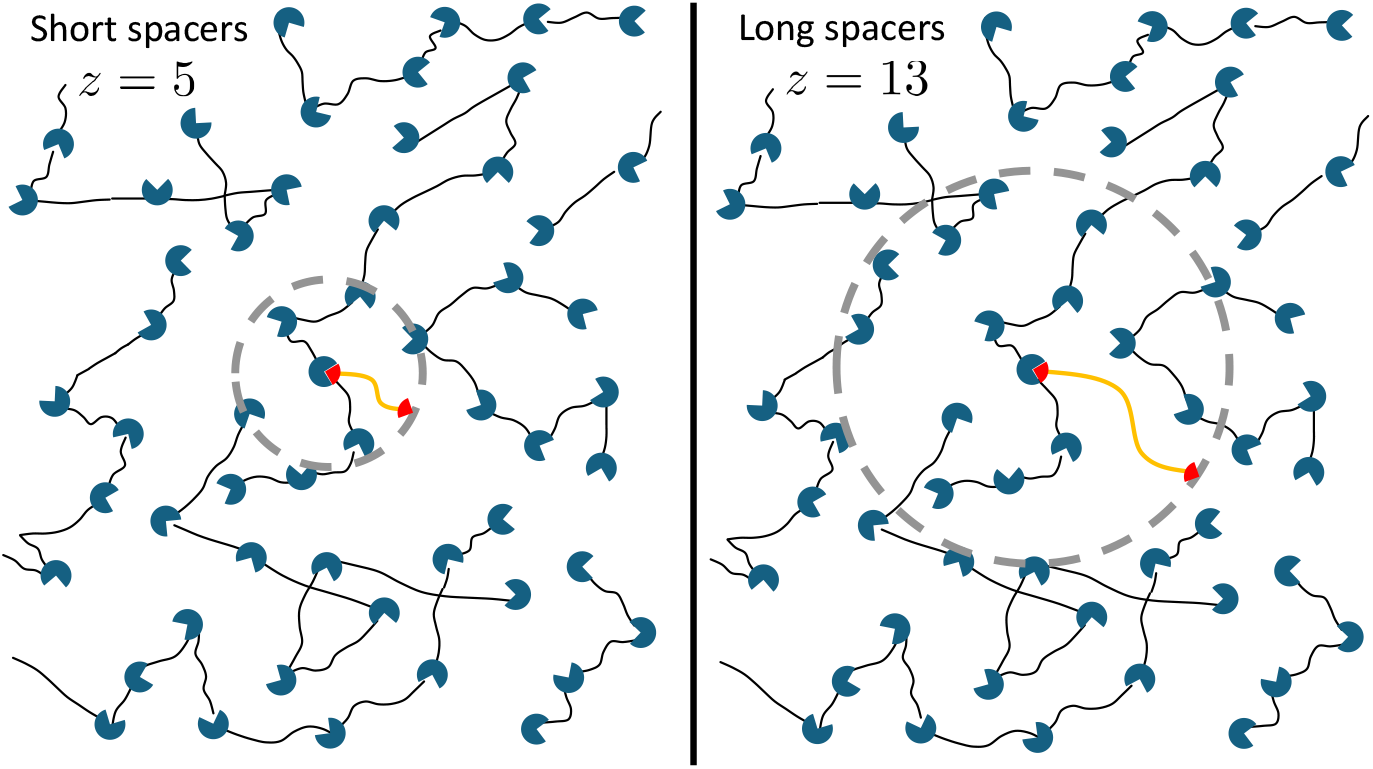
The B spacer (orange) limits the number of A stickers (blue) accessible to the unbound B sticker (red). (left) There are *z* = 5 A stickers within the dotted line, which indicates the volume within reach of the spacer. (right) A longer B spacer increases the accessible volume such that there are *z* = 13 stickers within the accessible volume.

Adjacent beads along each polymer backbone were connected using harmonic bond potentials,

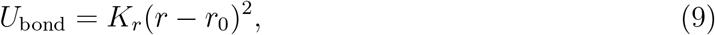

where *r*_0_ is the equilibrium bond distance (i.e., the bead diameter, 4.5 Å), *K*_*r*_ = 2 kcal*/*mol*/*Å^2^ is the bond constant, and *r* is the distance between two consecutive beads.

Angular rigidity was introduced via a cosine angle potential applied to all consecutive triplets of bonded beads, providing moderate chain stiffness,

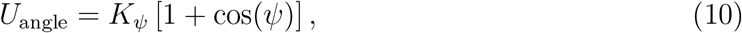

where *K*_*ψ*_ = 0.5 kcal*/*mol is the angle constant and *ψ* is the angle between two consecutive bonds.

Nonbonded interactions were modeled using truncated Lennard–Jones (LJ) potentials. Interactions between spacers and any other spacer/sticker beads were modeled as weakly attractive and characterized by an LJ energy scale corresponding to 0.3 *k*_B_*T*,

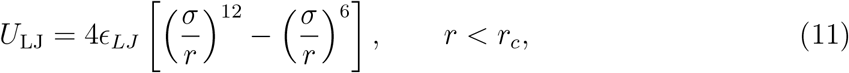

where *ϵ*_*LJ*_ is the LJ energy scale, *σ* = 4.5 Å is the bead diameter, *r* is the distance between two beads, and *r*_*c*_ = 11.25 Å is the cutoff distance.

Repulsive interactions between identical stickers (A–A and B–B) were modeled using shifted Lennard–Jones (Weeks–Chandler–Andersen-type) potentials to prevent unphysical aggregation and suppress hopping or bond-exchange artifacts,

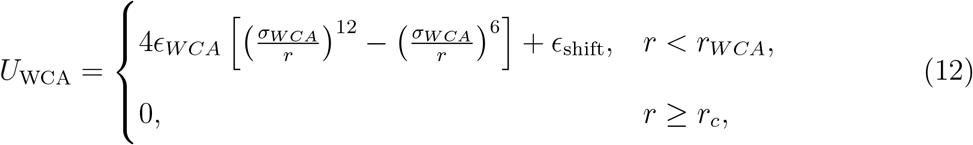

where *r*_*WCA*_ = 12.32 Å is the cutoff distance, *σ*_*WCA*_ is set to 11 Å, *ϵ*_*WCA*_ = 1*k*_*B*_*T*, and *ϵ*_shift_ shifts the LJ potential to zero at the cutoff, avoiding an energy discontinuity.

To enforce selective heterotypic association, strong specific sticker–sticker interactions were restricted exclusively to interactions between stickers of different types (i.e., between sticker A and sticker B). The strength of the A–B interaction was varied systematically from 2 to 5 *k*_B_*T* in increments of 0.5.

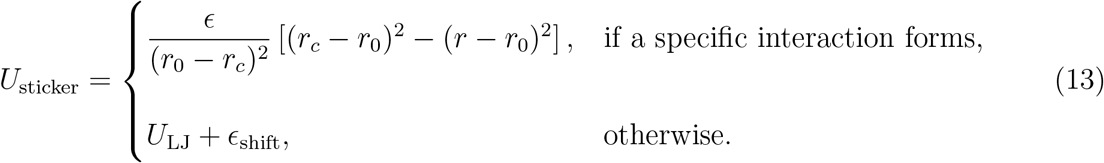

where *r*_0_ = 5.05 Å is the equilibrium interaction distance, *r*_*c*_ = 5.55 Å is the critical distance within which a specific interaction can form, and *ϵ*_shift_ shifts the LJ potential at the critical distance to avoid an energy discontinuity. The specific interaction energy has the value *ϵ* at *r* = *r*_0_.

Specific sticker–sticker associations were modeled dynamically using a combination of bond creation and bond breaking algorithms. When an A sticker and B sticker came within the critical distance *r*_*c*_, a probabilistic reaction formed a reversible bond, mimicking transient physical crosslinks. Bond breaking was allowed beyond the critical distance *r*_*c*_, ensuring dynamic equilibrium between bound and unbound sticker pairs. The parameters governing bond creation and breaking were chosen such that the system reached a steady-state distribution of sticker bonds within the simulation timescale.

The polymer volume fraction in the simulation box, defined as the fraction of volume occupied by polymer beads, was varied across simulations and set to 10%, 15%, 20%, or 25%. The simulation box size was adjusted accordingly while maintaining periodic boundary conditions in all three spatial dimensions (Fig. 3).

**Figure 3:**
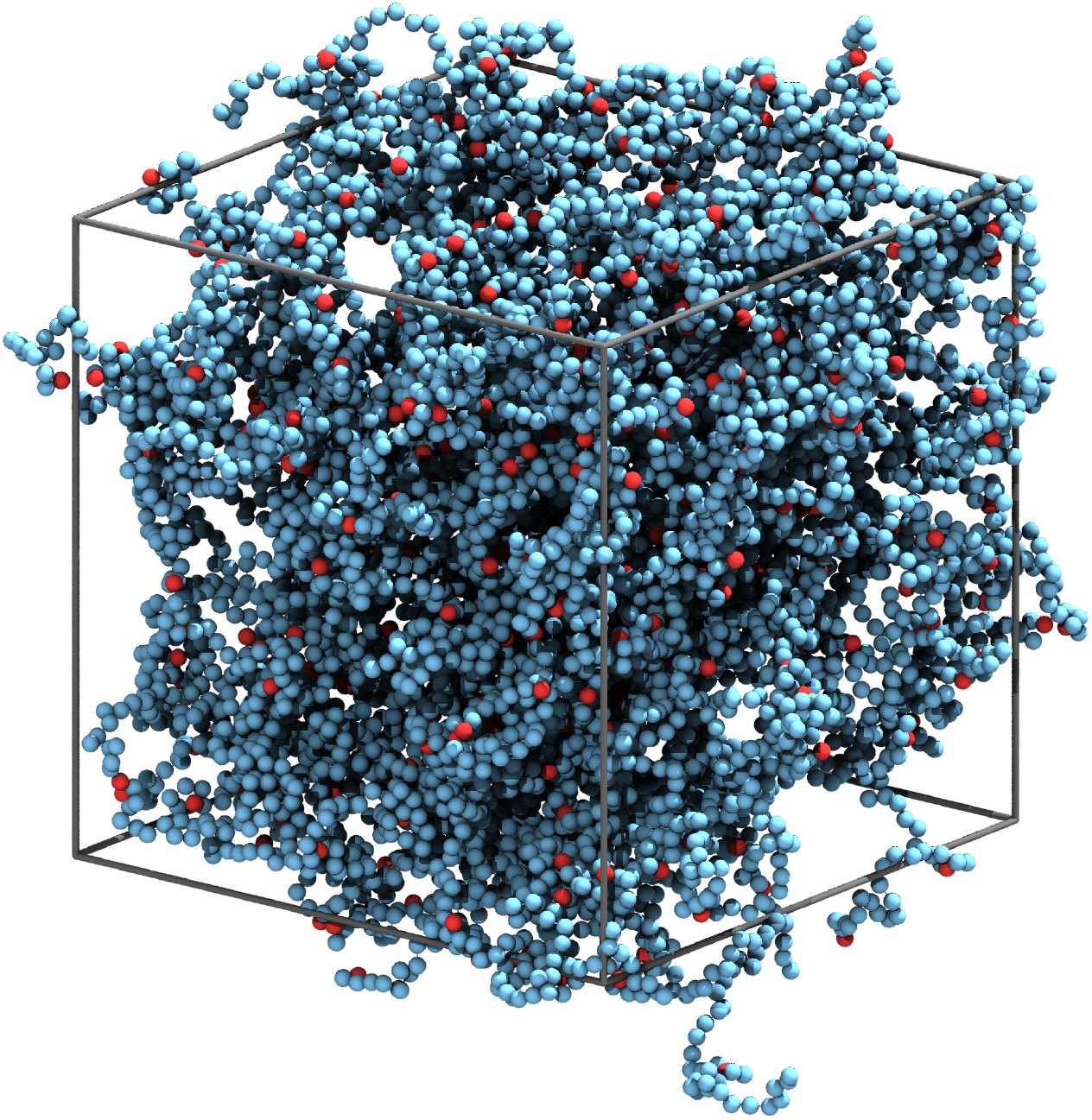
Representative simulation snapshot of sticker–spacer polymer chains. Stickers and spacers are shown as red and blue beads, respectively. The gray wireframe outlines the simulation box.

All simulations began with an energy minimization step to remove unfavorable overlaps and relax the initial configuration. Following minimization, an annealing protocol was applied to gradually introduce sticker–sticker bonding interactions. During annealing, the bond affinity associated with sticker interactions was ramped linearly from zero to its target value. This procedure prevented kinetic trapping and promoted equilibration of network topology.

Annealing simulations were performed in the NVT ensemble at a temperature of 310 K using a Langevin thermostat. The thermostat friction coefficient was chosen to reproduce the viscosity of water (10^*−*3^ Pa · s), thereby accounting for solvent effects through both viscous damping and stochastic forces.

After annealing, the final configuration was used as the starting point for long production runs. Production simulations were also carried out in the NVT ensemble at 310 K using the same Langevin thermostat parameters. No further annealing or parameter ramping was applied during this stage. To ensure proper sampling and avoid center-of-mass drift or inhomogeneous bead distributions, periodic load balancing was employed during long simulations. The equations of motion were integrated using a velocity–Verlet scheme with a timestep of 30 fs.

Production runs were carried out for up to 10^9^ timesteps, corresponding to 30 *µ*s of simulated time. For each simulation setup (i.e., volume fraction, sticker count, and sticker A–sticker B interaction strength), five independent trajectories were simulated with different random seeds. All reported observables were computed from steady-state segments of the production runs after verifying that global quantities such as energy and sticker bond number had converged.

### Comparison between theory and simulation

We used bond saturation for an initial comparison between our simulation and theory models. The number of bonds formed, *P*, can be computed by first applying the Stirling approximation to Eq. 8

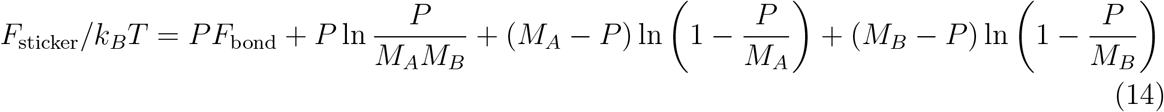

where 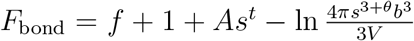 . Next, we minimize Eq. 14 with respect to *P*. The condition *∂F*_sticker_*/∂P* = 0 yields

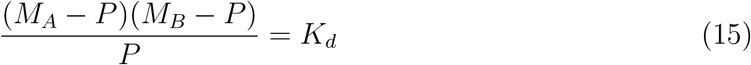

which shows that sticker bonding has the same form as a two component chemical equilibrium,^8,9,11^ where 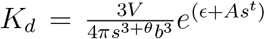 is the dissociation constant. Eq. 15 is a quadratic function with the solution

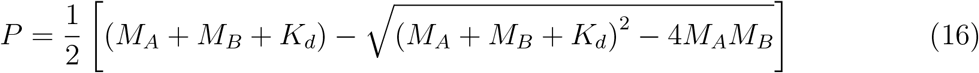

which can be further simplified in the case of symmetric mixtures *M*_*A*_ = *M*_*B*_ = *M*

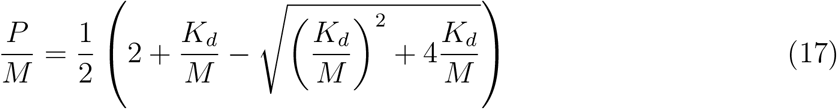

The ratio *K*_*d*_*/M*_*A*_ can be written in terms of the volume fraction *ϕ* = (*N*_*A*_ + *N*_*B*_)*Lπσ*^3^*/*(6*V* )

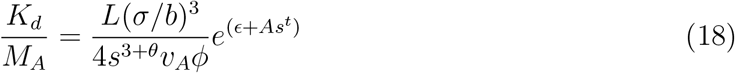

Fig. 4 compares Eq. 17 to the average bond saturation in our simulations. The optimal fit was obtained with *σ/b* = 0.78, a value consistent with the expectation that *b* is comparable to the Kuhn length, which is larger than the bead size. However, we cannot exclude the possibility that this parameter absorbs contributions from the unequal treatments of the linker as a distribution or uniform sphere in *ρ*(*R*) and *z*, respectively. Fig. 5 compares Eq. 16 to simulations with chains of unequal valence (*M*_*A*_/= *M*_*B*_), which confirms the observation that the theory is insensitive to spacers of unequal length. While the fit quality of Figs. 4 and 5 is encouraging, it must be emphasized that these calculations are equivalent to SR theory because the differences between our formulation and SR’s are absorbed by the fitting parameter.

**Figure 4:**
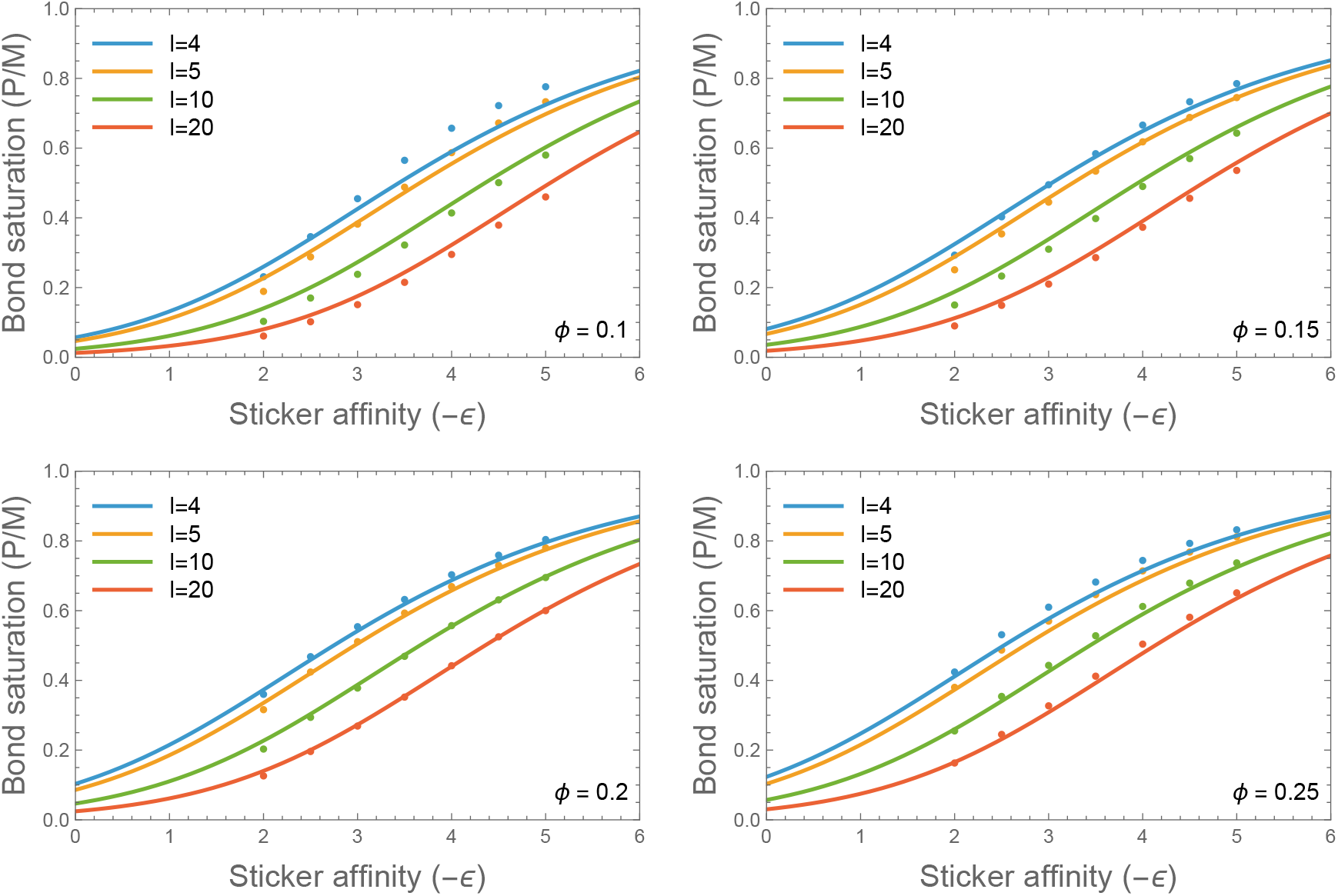
The predicted bond occupancy from Eq. 17 (lines) agrees well with simulations (points) conducted at fixed volume fraction *ϕ*, as indicated. The colors indicate chains of differing spacer length *l*, or equivalently, differing valence *v* = *L/l*. The single fitting parameter (*σ/b*)^3^ = 0.47 is the same for all curves. *n*_*A*_ = *n*_*B*_ = 50, *L* = 100

**Figure 5:**
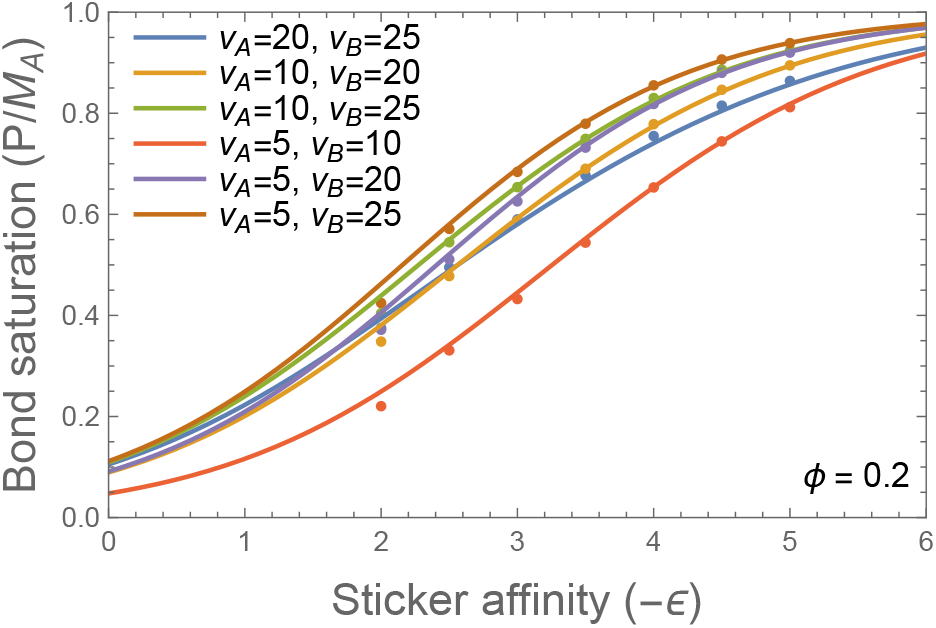
Comparison of the calculated bond saturation (Eq. 16) to simulations of chains with unequal valences. The fitting parameter, (*σ/b*)^3^ = 0.47, is unchanged from Fig. 4 *n*_*A*_ = *n*_*B*_ = 50, *L* = 100

The difference between our theory and SR lies in the coupling between the sticker and spacer degrees of freedom, which manifests in the spacer configurations. Fig. 6 compares the polymer end-to-end distribution function, Eq. 4, to the distribution of distances between adjacent stickers. To make this comparison, we obtain the characteristic length, *R*_0_, from the average extension measured in the simulation data 4*π ∫ρ*(*R*)*R*^3^*dR* = ⟨*R*⟩, which gives ⟨*R*⟩ = 0.954*R*_0_ for real chains.

**Figure 6:**
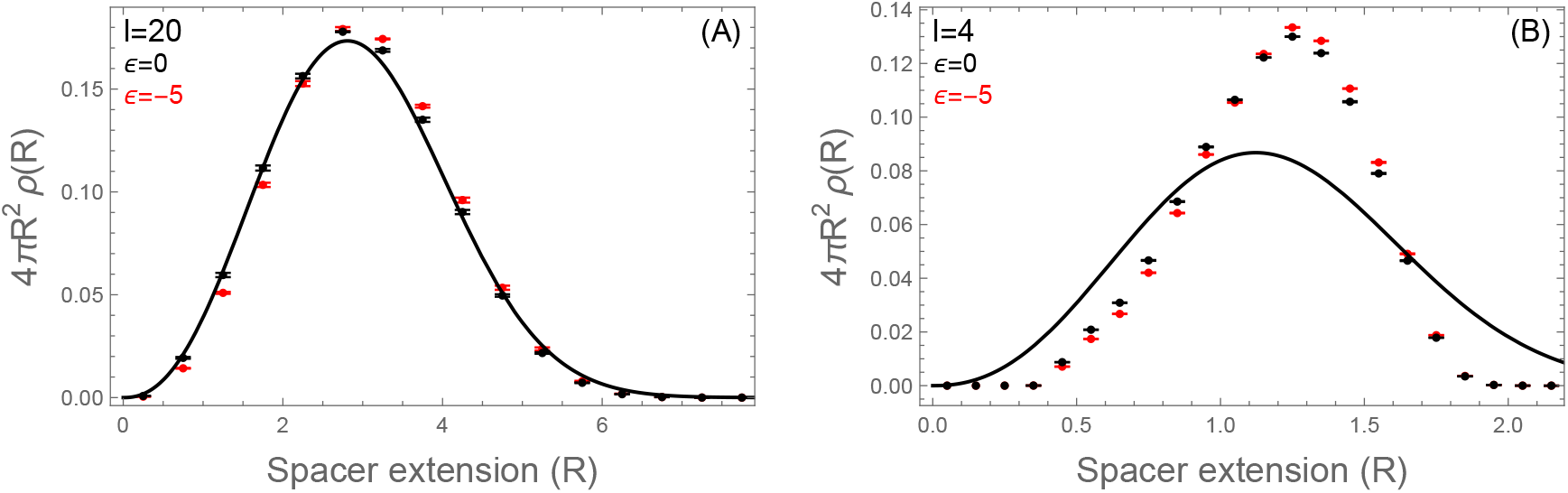
(A) The distribution of distances between adjacent stickers for chains with long spacers (*l* = 20) agrees well with the probability distribution function for a self-avoiding polymer. The black line shows 4*πR*^2^*ρ*(*R*) using the real chain values in Eq. 4. Increasing the affinity of the stickers from *ϵ* = 0 (black points) to *ϵ* = −5 (red points) causes a shift in the distribution toward more extended states. (B) The shift toward more extended states is also seen in chains with short spacers (*l* = 4), however, such short spacers are poorly described by Eq. 4.

For stickerless polymers (*ϵ* = 0, black points) the observed distribution is well-captured by Eq. 4. Increasing the sticker affinity to *ϵ* = −5 (Fig. 6, red points) leads to a subtle, but systematic, shift to more extended states. This shift is more obvious for shorter spacers, however, it is readily apparent that Eq. 4 is a poor description of the *l* = 4 segments shown in Fig. 6B. In particular, the observed distribution is more sharply peaked than the scaling limit described by Eq. 4, with a particularly steep cutoff at large extensions. This discrepancy arises in part from the chain rigidity introduced by the angular potential (Eq. 10), which, together with the finite contour length of short spacers, leads to deviations from the flexible-chain scaling statistics assumed in the theory.

To compare the observed spacer stretching to our theory, we must account for heterogeneity in the effective spacer length. Even in our simulations, where the stickers are uniformly spaced, the presence of non-bonded stickers will increase the effective spacer length for adjoining stickers. More precisely, a stretch of *n* − 1 unbound stickers will create an effective spacer of length *l*_eff_ = *nl*. Eq. 8 describes a stretching energy *As*^*t*^ per bond, implying that the stretching is distributed across *l*_**eff**_ . If *n* is small, as is usually the case in the condensed regime, a reasonable approximation is that the stretching is evenly distributed across the individual spacers. Therefore, each of the individual spacers in an effective spacer, will have an additional length (*s* − 1)*R*_*ϵ*=0_*/n*, where *R*_*ϵ*=0_ is the extension of an individual spacer in the absence of stickers, *ϵ* = 0. If there are *N*_*n*_ instances of effective spacers of length *nl*, then the average extension can be computed as

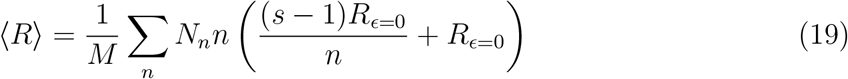

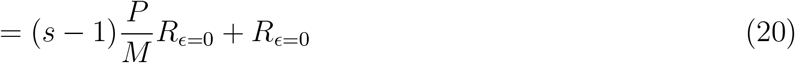

This formula predicts that the average spacer extension will increase with sticker affinity because more attractive bonds increase *P* . Fig. 7 compares the average distance between adjacent stickers to the predictions of Eq. 20. The calculation agrees favorably with simulations for long spacers, but deviates for the shorter spacers. This deviation can be understood from Fig. 6B, which shows that short spacers are more constrained in long extension states than what is described by *ρ*(*R*) (Eq. 4). The abrupt cutoff at large *R* indicates stiffer effective spring constant, which is consistent with the simulation spacers being less extended than the prediction of Eq. 20.

**Figure 7:**
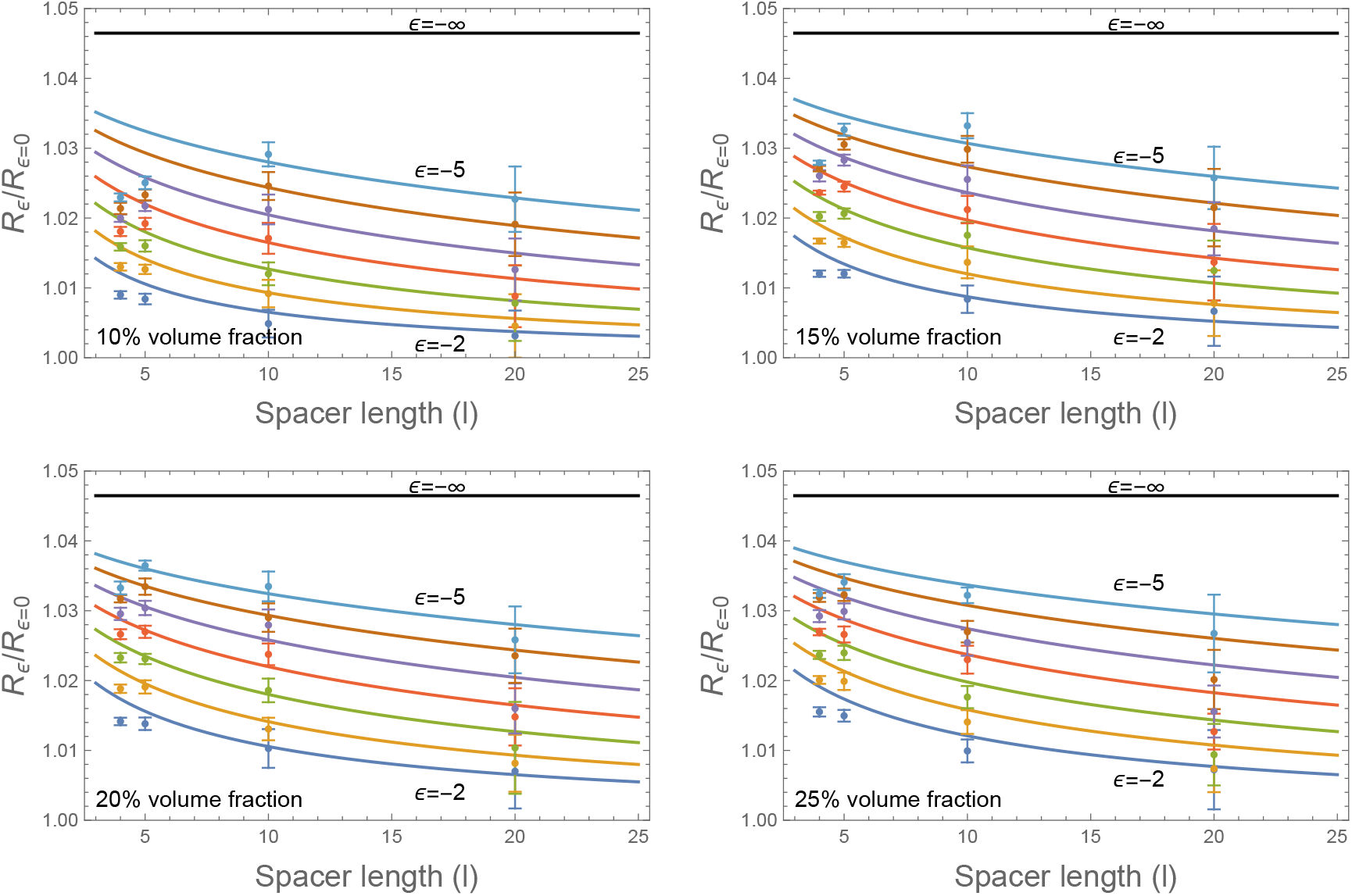
The average sticker-sticker distance *R* increases with binding affinity *ϵ*. The plot shows the ratio of sticker-sticker distances for chains with attractive stickers *R*_*ϵ*_ to chains with inert stickers *R*_*ϵ*=0_. The observed stretching is consistent with a weighted average of stretched spacers and unstretched spacers (Eq. 20 with Eq. 17).

## Discussion

We set out to understand how the physical connections between adjacent stickers influences the sticker binding free energy. We found that connectivity contributes to the free energy via the loss of spacer conformational entropy and a gain in the number of sticker-sticker combinations that can form. The magnitude of these terms can be estimated by ln *z*, suggesting that they are each on the order of a few *k*_*B*_*T* per bond. However, because these factors oppose each other there is a substantial cancellation and, in fact, the dependence on the spacer length vanishes. The cancellation of length-dependent terms allows the stickers to behave like untethered reactants, thereby validating the SR approximation in which the sticker free energy depends only on the sticker concentration. Fortuitously, the spacer term that remains after the cancellation, which describes the elastic stretching of spacers, is length-independent and can be absorbed into either the bond energy *ϵ* or the proximity factor *v*_0_*/V* . Interestingly, the spacer stretching result does not emerge from constructing the partition function of a sticker-spacer unit within a uniform background of binding sites. This is because the gain of binding partners has the same *R*^2^*dR* scaling as the conformational phase space of *ρ*(*R*). Instead, the stretching reflects a competition between the *local* entropy of a spacer and the *global* entropy of bonding combinatorics (Eq. 2).

Further insight into the irrelevant of explicit connections on the sticker binding statistics can be obtained from the correspondence between Eq. 15, and the equilibrium of two-component reactions. In chemical equilibrium the extent of the reaction depends on the concentration of reactants but not on the size of the reaction volume. Therefore, we can interpret the spacer connectivity effect as determining the “container” that confines a particular sticker.^18^ Increasing the length of a particular spacer increases the search volume, but does not change the concentration of binding partners. Thus, the formation of a sticker-sticker bond incurs the same tradeoff between binding energy and translational entropy as the formation of heterodimers between untethered reactants.

Our formulation of the sticker-spacer theory provides clarity as to when connectivity can be neglected and when it cannot. For instance, natural biopolymers often have significant heterogeneity, which can take the form a variety of spacer lengths. Our theory confirms that heterogeneity of spacer lengths can be handled within the SR mean-field approximation, thereby providing considerable complexity reduction. ^19^ This conclusion assumes that the spacers remain well-mixed, which is ensured when the spacers are part of the same molecule.[Akram et al., in preparation] Similarly, a difference in spacer lengths between the A and B molecules is irrelevant since the spacer cancellation restores symmetry between the A and B molecules. An extreme case of spacer asymmetry is when one scaffold is a multivalent globular protein such as SPOP or EPYC.^12,20^ Our theory shows that this situation can still be described within the SR formalism using the mean-field concentration of binding sites, although care should be taken to account for the perturbation the globular domain imposes on the spacer entropy. ^21^

Our theory also helps to anticipate situations where the connectivity constraint may induce deviations from mean-field behavior. The irrelevance of connectivity in SR theory occurs because the loss of conformational entropy and the gain in potential binding partners both scale with the volume accessible to the spacer. Therefore, connectivity could be expected to re-enter the theory in cases where this equivalence is broken. For example, if there are significant heterogeneities in the dense phase concentration, then it can be more advantageous for a bonds to form consecutive bonds between the same molecules. ^11,22^ Another such situation occurs at the surface of the dense phase. This is because a spacer near the interface does not gain binding partners by extending outside the dense phase but still incurs the full conformational penalty upon binding. Similarly, when oligomers form in the dilute phase, increasing spacer lengths do not result in an increase in potential binding partners. As a result the conformational entropy loss in the dilute phase can be deterministic in the propensity for condensation. ^13^ The present formulation assumes that individual stickers form at most one bond and that stickers and spacers have comparable steric length scales. Allowing multiple simultaneous bonds per sticker or substantial sticker–spacer size asymmetry could modify the combinatorial and conformational entropy contributions, potentially introducing corrections beyond a simple renormalization of the effective binding affinity and will be explored elsewhere.

## Conclusion

We have reformulated the two-fluid model for associative polymers to account for the interplay between spacer conformations and sticker binding. We find that mean-field models that ignore the relationship between these factors are successful because of a fortuitous cancellation between two terms that depend on spacer length. Our theory shows how the combinatorial entropy of bond formation distorts the conformations of spacers and provides insights into situations, like interfaces, where the conformational entropy of spacers is likely to have contributions to the free energy that differ from the mean-field picture.

## Acknowledgements

This work was supported by NIH grant R01GM141235. The author(s) would like to thank the Isaac Newton Institute for Mathematical Sciences, Cambridge, for support and hospitality during the programme SPL03, where work on this paper was undertaken. This research was supported in part by grant NSF PHY-2309135 and the Gordon and Betty Moore Foundation Grant No. 2919.02 to the Kavli Institute for Theoretical Physics (KITP).

